## Supplementary information for "Unsupervised alignment reveals structural commonalities and differences in neural representations of natural scenes across individuals and brain areas"

*Description of the additional dataset: Natural movie 1 and Natural movie 3*

In the main text, we analyzed the neural representations of natural scenes. Here, we provide the results of the same analysis using two additional datasets: Natural movie 1 and Natural movie 3. Natural movie 1 consists of a 30-second continuous movie stimulus, and Natural movie 3 consists of a 120-second continuous movie stimulus. We segmented them into 1/3-second short movie stimuli and treated them as 90 segmented movie stimuli for Natural movie 1 and 360 segmented movie stimuli for Natural movie 3. There were a total of 32 mice for the experiment of natural scenes stimuli, and all of them were used in the analysis.

After the standardization of the spike counts for each neuron, we then computed the trial average of spike counts during each 1/3-second short movie stimulus and considered these as the neural responses of the pseudo-mice to be aligned.

As in the main text, we analyzed the following 8 areas in the visual cortex and subcortical regions: 6 area of the visual cortex (VISp: primary visual area, VISrl: rostrolateral visual area, VISl: lateromedial visual area, VISal: anterolateral visual area, VISpm: posteromedial visual area: VISam:

*Corresponding author
[1] These authors contributed equally to this work

anteromedial visual area), 1 area of the thalamus (LGd: dorsal part of the lateral geniculate nucleus), and 1 area of the hippocampus (CA1: cornu ammonis 1). The 6 areas in the visual cortex and LGd are components of the visual system, while CA1 is a part of the memory system. Not all brain areas were measured in each mouse, and the number of neurons used for analysis ranged from 20 to 200 per mouse per region, with an average of approximately 60.

To obtain a statistically reliable estimate of representational similarity structures, we group-averaged the neural activity of multiple mice, considered as a "pseudo-mouse". To construct a pair of pseudo-mice, we randomly divided the total of 32 mice provided in the dataset into two non-overlapping groups of 16 mice each. We then concatenated the neural representations within each group along the neurons. Therefore, the neural representations for each pseudo-mouse are represented by vectors of 400 to 1,200 neurons, varying by brain area. Using these aggregated neural representations, we estimated the group-averaged representational similarity structures for the 90 or 360 segmented movie stimuli in each area for each pseudo-mouse. The dissimilarity between two stimuli was quantified using cosine distance, based on the trial-averaged normalized spike counts.

The procedures of the analysis were the same as in the main text. Below, we show the results of the analysis.

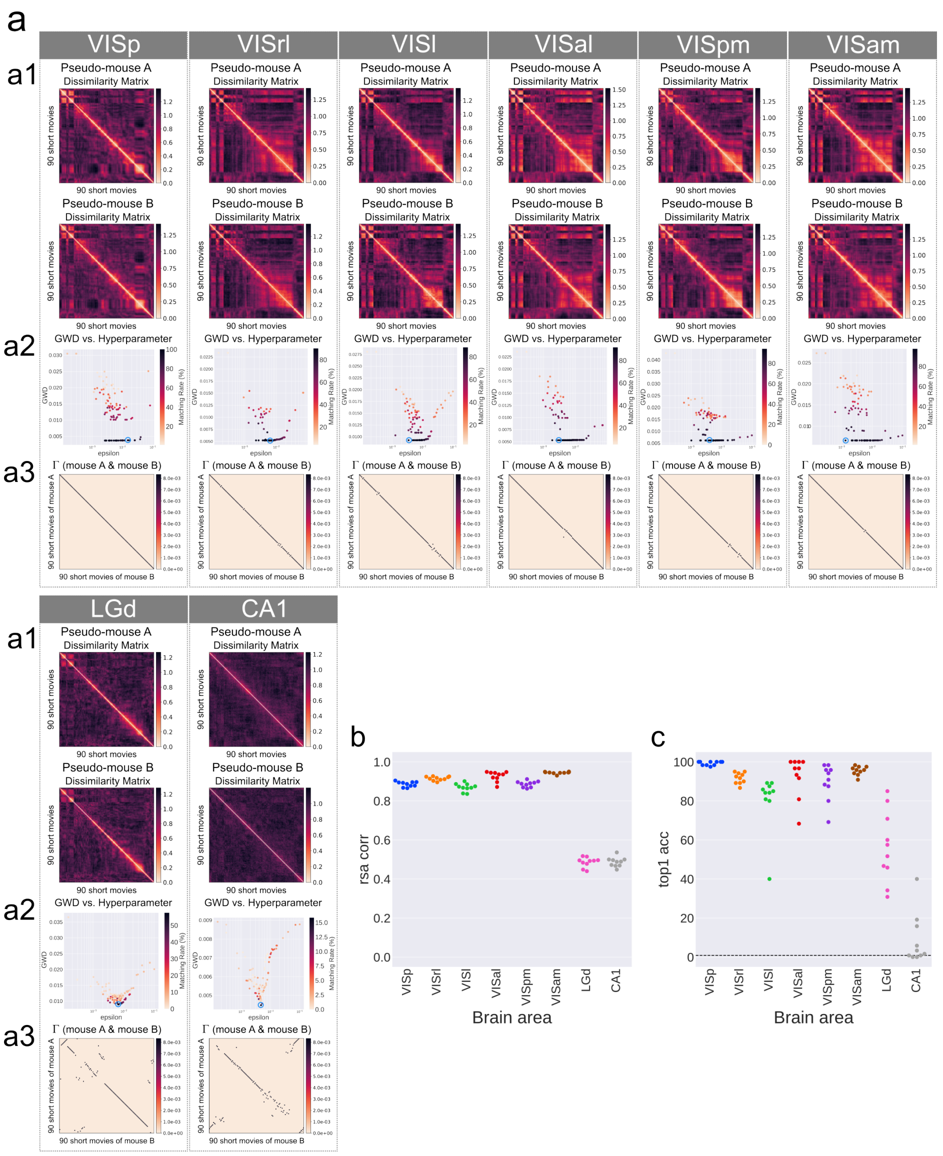


Figure S1: **Unsupervised alignment between the same areas in different pseudo-mice for Natural movie 1** (a) Results of GWOT between the dissimilarity matrices of the same areas in different pseudo-mice. (a1) Dissimilarity matrices of a pair of pseudo-mice for the 90 stimuli in each area. (a2) Relationship between GWD (objective of GWOT) and the hyperparameter $\epsilon$. (a3) Optimal transportation plan $\Gamma^*$ between the dissimilarity matrices of a pair of pseudo-mice. This matrix corresponds to the point encircled in blue in Fig. S1a2. (b) Correlation coefficient of RSA between the dissimilarity matrices of a pair of pseudo-mice across 10 trials. (c) Top 1 matching rate of the unsupervised alignment between the dissimilarity matrices of a pair of pseudo-mice across 10 trials.

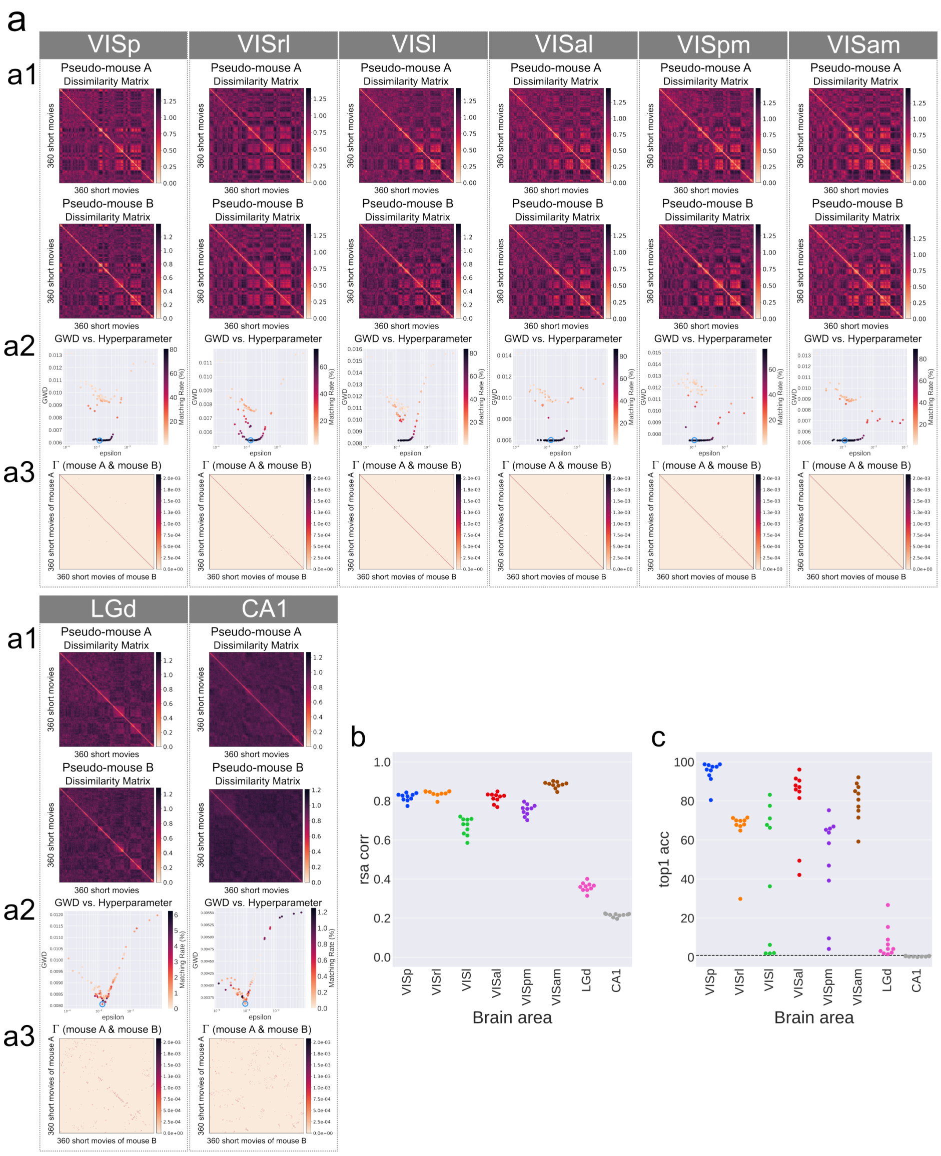


Figure S2: **Unsupervised alignment between the same areas in different pseudo-mice for Natural movie 3** (a) Results of GWOT between the dissimilarity matrices of the same areas in different pseudo-mice. (a1) Dissimilarity matrices of a pair of pseudo-mice for the 360 stimuli in each area. (a2) Relationship between GWD (objective of GWOT) and the hyperparameter $\epsilon$. (a3) Optimal transportation plan $\Gamma^*$ between the dissimilarity matrices of a pair of pseudo-mice. This matrix corresponds to the point encircled in blue in Fig. S3a2. (b) Correlation coefficient of RSA between the dissimilarity matrices of a pair of pseudo-mice across 10 trials. (c) Top 1 matching rate of the unsupervised alignment between the dissimilarity matrices of a pair of pseudo-mice across 10 trials.

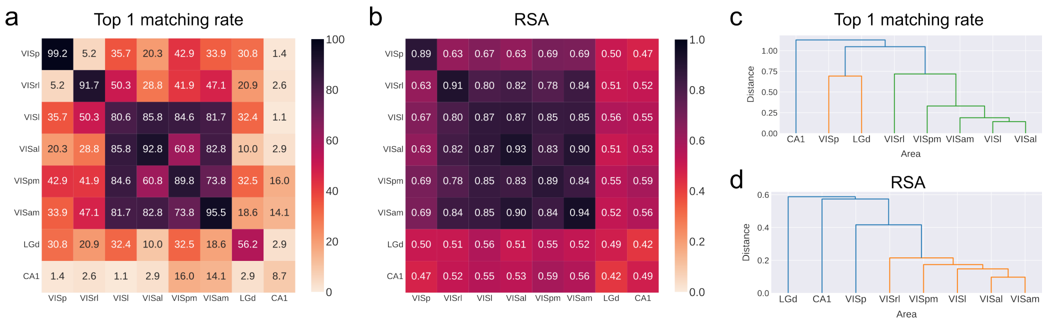


Figure S3: **Unsupervised alignment between the different areas in different pseudo-mice for Natural movie 1** (a) Average top 1 matching rate of the unsupervised alignment for each pair of brain areas. (b) Average correlation coefficient of the RSA for each pair of brain areas. (c) Hierarchical clustering of brain areas based on the average top 1 matching rate. Here, the distance between areas is defined as $(100 - \text{top 1 matching rate})/100$ and Ward's method is employed as the clustering criterion. (d) Hierarchical clustering of brain areas based on the average correlation coefficient. The distance between areas is defined as $(1 - \text{correlation})$.

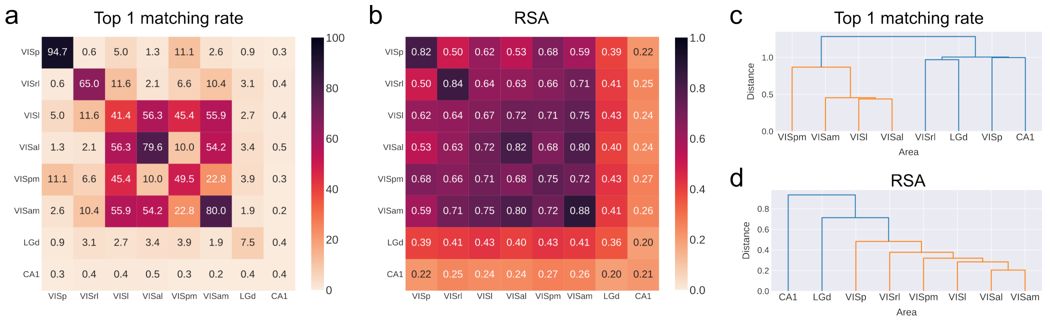


Figure S4: **Unsupervised alignment between the different areas in different pseudo-mice for Natural movie 3** (a) Average top 1 matching rate of the unsupervised alignment for each pair of brain areas. (b) Average correlation coefficient of the RSA for each pair of brain areas. (c) Hierarchical clustering of brain areas based on the average top 1 matching rate. Here, the distance between areas is defined as $(100 - \text{top 1 matching rate})/100$ and Ward's method is employed as the clustering criterion. (d) Hierarchical clustering of brain areas based on the average correlation coefficient. The distance between areas is defined as $(1 - \text{correlation})$.
